# Crystal structure of FAD-independent methylene-tetrahydrofolate reductase from *Mycobacterium hassiacum*

**DOI:** 10.1101/2022.11.18.516900

**Authors:** Manuel Gehl, Ulrike Demmer, Ulrich Ermler, Seigo Shima

## Abstract

FAD-independent methylene-tetrahydrofolate (methylene-H_4_F) reductase (Mfr), recently identified in mycobacteria, catalyzes the reduction of methylene-H_4_F to methyl-H_4_F with NADH as hydride donor by a ternary complex mechanism. This biochemical reaction corresponds to that of the ubiquitous FAD-dependent methylene-H_4_F reductase (MTHFR), although the latter uses a ping-pong mechanism with FAD as prosthetic group. Comparative genomics and genetic analyses indicated that Mfr is indispensable for the growth of *Mycobacterium tuberculosis*, which lacks the MTHFR-encoding gene. Thus, Mfr is an excellent target enzyme for the design of antimycobacterial drugs. Here, we report the heterologous production, enzymological characterization and the crystal structure of Mfr from the thermophilic mycobacterium *M. hassiacum* (hMfr), which shows 78% sequence identity to Mfr from *M. tuberculosis.* Although hMfr and MTHFR show very low sequence identity and different catalytic mechanisms, their tertiary structures are highly similar, which suggests a divergent evolution of Mfr and MTHFR from a common ancestor. Most of the important active-site residues of MTHFR are conserved and equivalently positioned in the tertiary structure of hMfr. The Glu9Gln variant of hMfr exhibits a drastic reduction of the catalytic activity, which supports the predicted function of the glutamate residue as proton donor in both Mfr and MTHFR. The predicted nicotinamide binding site of hMfr is substantially narrower than the isoalloxazine binding site of MTHFR, which may reflect an evolutional adaptation to the different sizes of the coenzymes.

## 1 INTRODUCTION

Estimated ten million people are infected with tuberculosis each year and more than one million succumb to the desease.^1^ Enhanced by increasing drug resistance, tuberculosis poses an outstanding public health threat.^2^ Therefore, the development of novel antimycobacterial agents is urgently needed.^3^ One of the key challenges in antimicrobial drug development is finding new enzyme targets that are unique to the pathogen.^4, 5^

Methylene-tetrahydrofolate (methylene-H_4_F) reductase (MTHFR) is a ubiquitous enzyme involved in the central carbon metabolism of eukaryotes, bacteria and most archaea. This enzyme catalyzes the reduction of methylene-H_4_F to methyl-H_4_F using NAD(P)H as reducing agent.^6^ MTHFR of *E.coli* is a homotetramer of 33 kDa per monomer and contains FAD as prosthetic group. Catalysis proceeds via a ping-pong mechanism consisting of two half reactions. In the reductive half reaction, FAD is reduced by NAD(P)H and the generated FADH2 subsequently reduces methylene-H_4_F in the oxidative half reaction. The structure of the inactive variant of MTHFR from *E. coli* complexed with methyl-H_4_F strongly contributed to establish a catalytic scenario in combination with mutational kinetic analyses.^7–11^ In addition, crystal structures of MTHFR from *Thermus thermophilus, Saccharomyces cerevisiae* and *Homo sapiens* were reported.^12–15^

In the majority of the genome of mycobacteria, a canonical MTHFR gene was not found. Young et al. predicted by comparative genomic analysis and structure modeling that a gene of *M tuberculosis* may encode a non-canonical MTHFR.^16^ Later, Sah et al. found that the enzymes encoded by MSMEG_6596 and MSMEG_6649 in *Mycobacterium smegmatis* are non-canonical MTHFRs that do not contain FAD or any other prosthetic group. Nevertheless, those enzymes have methylene-H_4_F reductase activity (Figure 1). The amino acid sequence identity of the canonical and non-canonical MTHFRs was very limited, in which only 16-17% of sequence identity and a coverage of 13-26%.were found. The deletion of the MSMEG_6596 locus of *M. smegmatis* rendered the strain partially auxotrophic for methionine.^17^ Recently, Yu et al. reported that the protein encoded by Rv2172c of *M. tuberculosis* exhibits the same properties as the non-canonical MTHFR found in *M smegmatis*. Gene deletion experiments revealed the Rv2172c-locus is essential for the growth of *M. tuberculosis*.^18^ Apparently, the non-canonical MTHFR is the only enzyme catalyzing the reduction of methylene-H_4_F in *M. tuberculosis.* Since the non-canonical MTHFR found in mycobacteria is a new type of methylene-H_4_F reductase,^17^ this enzyme is referred to as **m**ethylene-tetrahydro**f**olate **r**eductase (Mfr) in this paper to distinguish it from canonical MTHFR.

**Figure 1:**
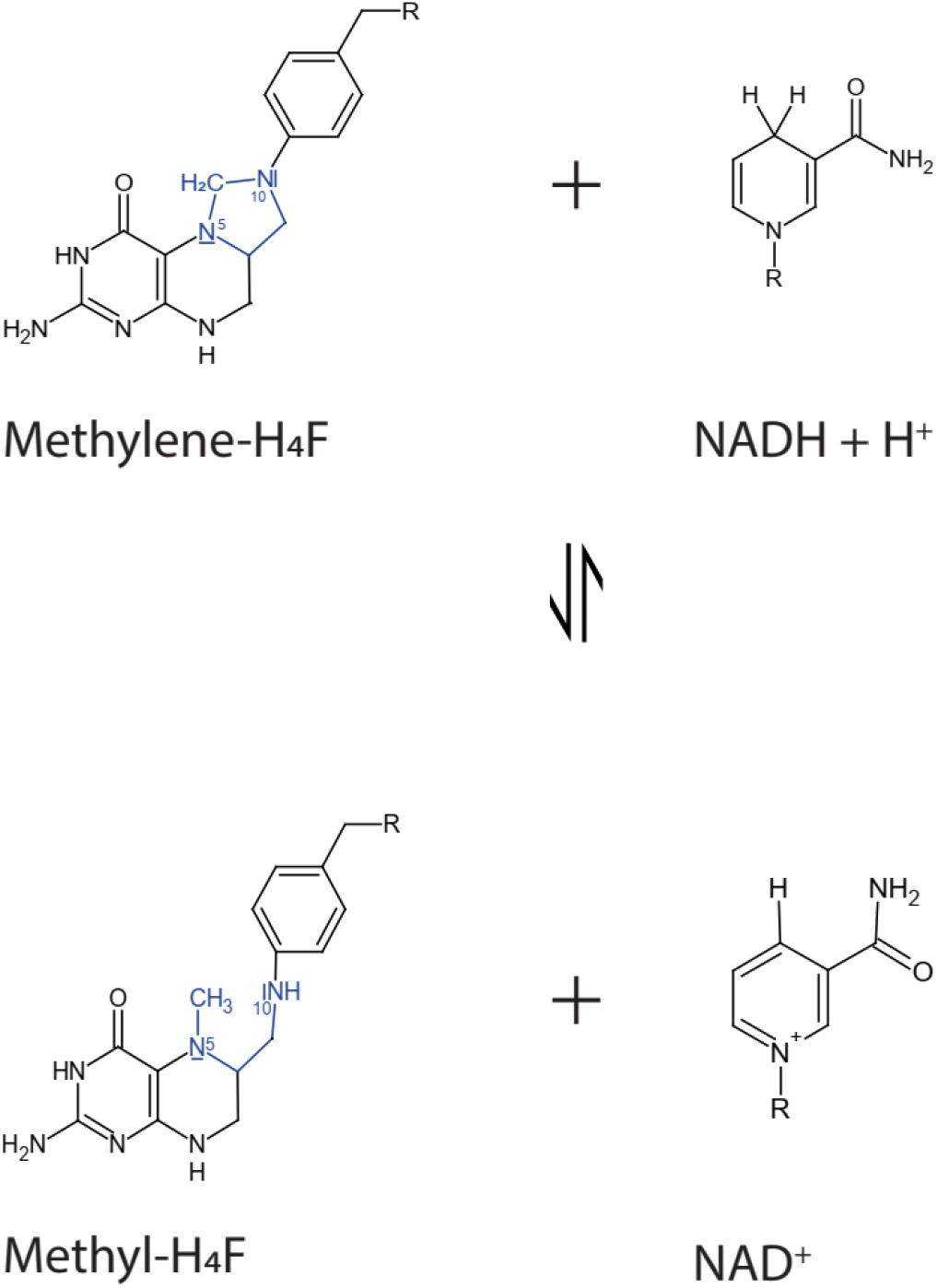
The reaction catalyzed by hMfr. A hydride is directly transferred from NADH to the carbon 14a position of methylene-H_4_F that is part of the imidazolidine ring (blue).

Since the genes encoding Mfr are found only in mycobacteria and some related bacteria, Mfr is an excellent target enzyme for the design of antimycobacterial agents. For designing Mfr inhibitors, a high-resolution structure of Mfr is required. Here, we present the crystal structure of Mfr from *Mycobacterium hassiacum* (hMfr). *M. hassiacum* is a thermophilic *Mycobacterium*, which is able to grow at 40–65 °C.^19^ The thermophilic properties of the enzymes from *M. hassiacum* are advantageous for crystallization.^20^ In addition, hMfr has 78% sequence identity (query coverage 87%) with its homolog from *M.* tuberculosis. Pronounced structural similarities between the active sites of hMfr and MTHFR suggested a common catalytic mechanism corresponding to the oxidative half reaction of MTHFR. To test this hypothesis, we performed a mutation analysis of the functional glutamate residue of Mfr.

## 2. RESULTS AND DISCUSSION

### 2.1 Purification and characterization of hMfr

The hMfr encoding gene was codon-optimized for the expression in *E. coli*, synthesized and cloned into pET-24b(+). The heterologously over-produced protein was purified to homogeneity (Table 1 and Figure 2A). The UV-visible spectrum of hMfr revealed only one peak at 280 nm, which confirms the absence of FAD (Figure 2B) as reported for Mfr from *M. smegmatis* and *M. tuberclosis*. ^17^, ^18^ For crystallization, the purified enzyme fraction was further purified by size exclusion chromatography. hMfr was eluted as a single symmetric peak at an elution volume corresponding to a molecular mass of approximately 30 kDa (Figure 2C), which indicates that the purified hMfr is in a monomeric form because the genome-deduced molecular mass of hMfr is 32 kDa. The apparent *V*_max_ and *K*_m_ values of hMfr are 18 μmole min^−1^mg^−1^ and 160 μM for methylene-H_4_F, respectively (Table 2 and Figure S1). The biophysical and enzymatic properties of hMfr are in accordance with the those previously published Mfr from *M. smegmatis* (see Table 2).^17^ hMfr shows an optimum temperature at 70 °C (Figure 3A). Thermostability of hMfr was evaluated by measuring enzymatic activity after incubation of the crude extract from *E. coli* at certain temperatures for 20 min. hMfr did not lose its catalytic competence until 50 °C (Figure 3B).

**Table 1:** Purification of hMfr expressed in *E. coli.* Cell extract was prepared from 3 g of cells (wet mass) and the enzyme was purified aerobically as described in the materials and methods section. One unit (U) activity refers to the oxidation of 1 μmole NADH per min to form methyl-H_4_F from methylene-H_4_F.

|  | Activity<br>(U) | Protein<br>(mg) | Specific<br>Activity<br>(U/mg) | Yield<br>(%) | Purification<br>Factor (fold) |
| --- | --- | --- | --- | --- | --- |
| Cell extract | 1200 | 200 | 6 | 100 | 1.0 |
| 50 °C heat treatment | 1000 | 100 | 10 | 82 | 1.6 |
| 40% ammonium sulfate | 900 | 86 | 10 | 73 | 1.7 |
| Phenyl-Sepharose HP | 500 | 49 | 10 | 43 | 1.7 |
| Resource Q | 300 | 19 | 16 | 25 | 2.6 |

**Figure 2:**
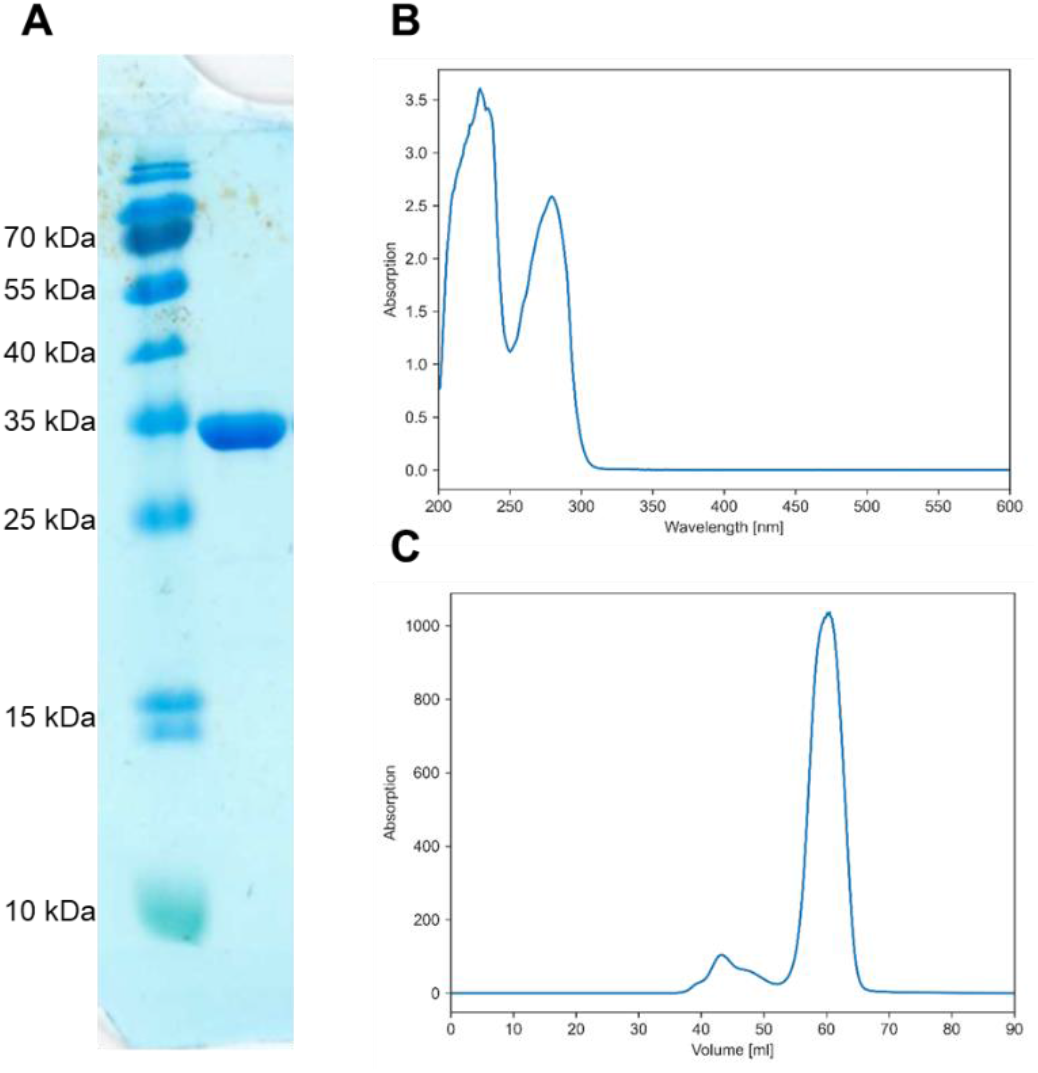
Characterization of hMfr purified from recombinant *E. coli.* (A) SDS-PAGE of purified hMfr (see also Table 1). (B) UV/Vis spectrum of purified hMfr (2.5 mg/ml) in 100 mM MOPS/NaOH pH 7.0 using a 1 cm light-path quartz cuvette. (C) Size-exclusion chromatogram of the purified hMfr preparation using a HiPrep 26/60 Sephacryl S-100 HR column.

**Table 2:** Kinetic properties of the hMfr wild type, hMfr E9Q mutant and Mfr proteins from *M. smegmatis* (MSMEG_6596 and MSMEG_6649). The data of Mfr from *M. smegmatis* were obtained from the previous publication.^17^

| Enzyme | Substrate | app$K_m$<br>(μM) | app$V_{max}$ (μmole<br>$\text{min}^{-1}\text{mg}^{-1}$) | app$k_{cat}$($\text{min}^{-1}$) | app$k_{cat}/\text{app}K_m$<br>($\text{min}^{-1}\mu\text{mole}^{-1}$) |
| --- | --- | --- | --- | --- | --- |
| hMfr wild | Methylene-<br>H<sub>4</sub>F | 160 ± 62 | 18 ± 2.6 | 590 ± 85 | 3.9 ± 1 |
| | NADH | 16 ± 3 | 17 ± 0.7 | 540 ± 23 | 35 ± 5 |
| hMfr E9Q | Methylene-<br>H<sub>4</sub>F | 250 ± 59 | 0.041 ± 0.004 | 1.3 ± 0.1 | 0.005 ± 0.0007 |
| | NADH | 44 ± 13 | 0.035 ± 0.004 | 1.1 ± 0.1 | 0.027 ± 0.054 |
| MSMEG6596 | Methylene-<br>H<sub>4</sub>F | 63 ± 11 | 7 ± 0.3 | 240 ± 10 | 3.8 ± 1 |
| | NADH | 33 ± 8 | 11 ± 1 | 370 ± 31 | 11 ± 4 |
| MSMEG6649 | Methylene-<br>H<sub>4</sub>F | 150 ± 30 | 1.2 ± 0.1 | 41 ± 3 | 0.28 ± 0.1 |
| | NADH | 110 ± 30 | 2.3 ± 0.3 | 80 ± 10 | 0.73 ± 0.3 |

**Figure 3:**
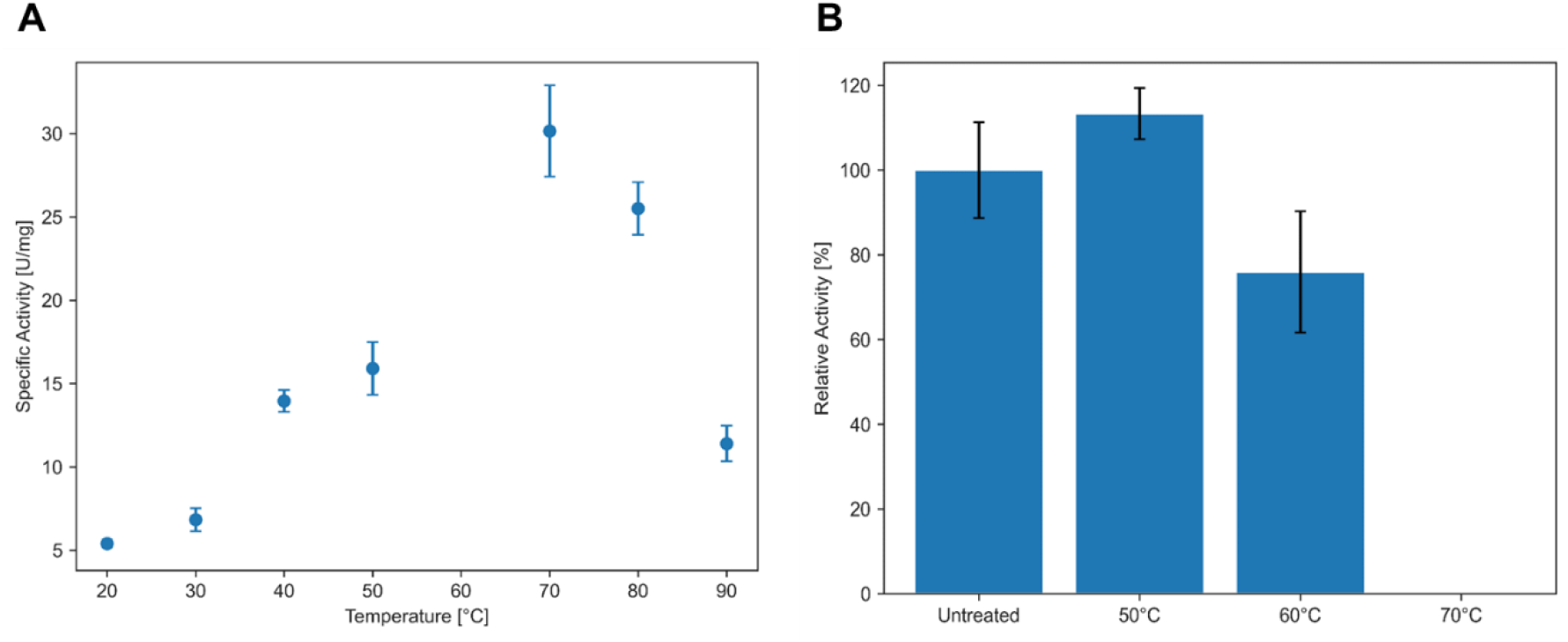
Thermophilic properties of hMfr. (A) Temperature dependency of the activity. (B) Residual activity in the cell extract after heat treatment for 20 min. Every data point consists of three independent measurements. The error bars represent the corresponding standard deviation.

### 2.2 Crystal structure of hMfr

Purified hMfr was crystallized by the sitting-drop vapor-diffusion method. The crystal of hMfr contains one monomer in the asymmetric unit and diffracted to 1.8 Å resolution (Table 3). The phase problem was solved by the molecular replacement method using a model structure generated by Alphafold2 as template (Figure S2). The core structure of this hMfr represents an β_8_α,_8_ or triose-phosphate isomerase (TIM) barrel fold. As normally found in TIM barrel family enzymes, several prolonged and specifically shaped structural segments between the secondary structure elements play an essential functional role. In hMfr, these inserted segments form the upper part of a putative active-site cleft on the C-terminal end of the β_8_α_8_ barrel (Figure 4). The α-helix bundle (212-243) after β7 and two elongated loops (loop 1:51-69 after β2 and loop 2: 118-134 after β4) form the outer wall of the active site cleft.

**Figure 4:**
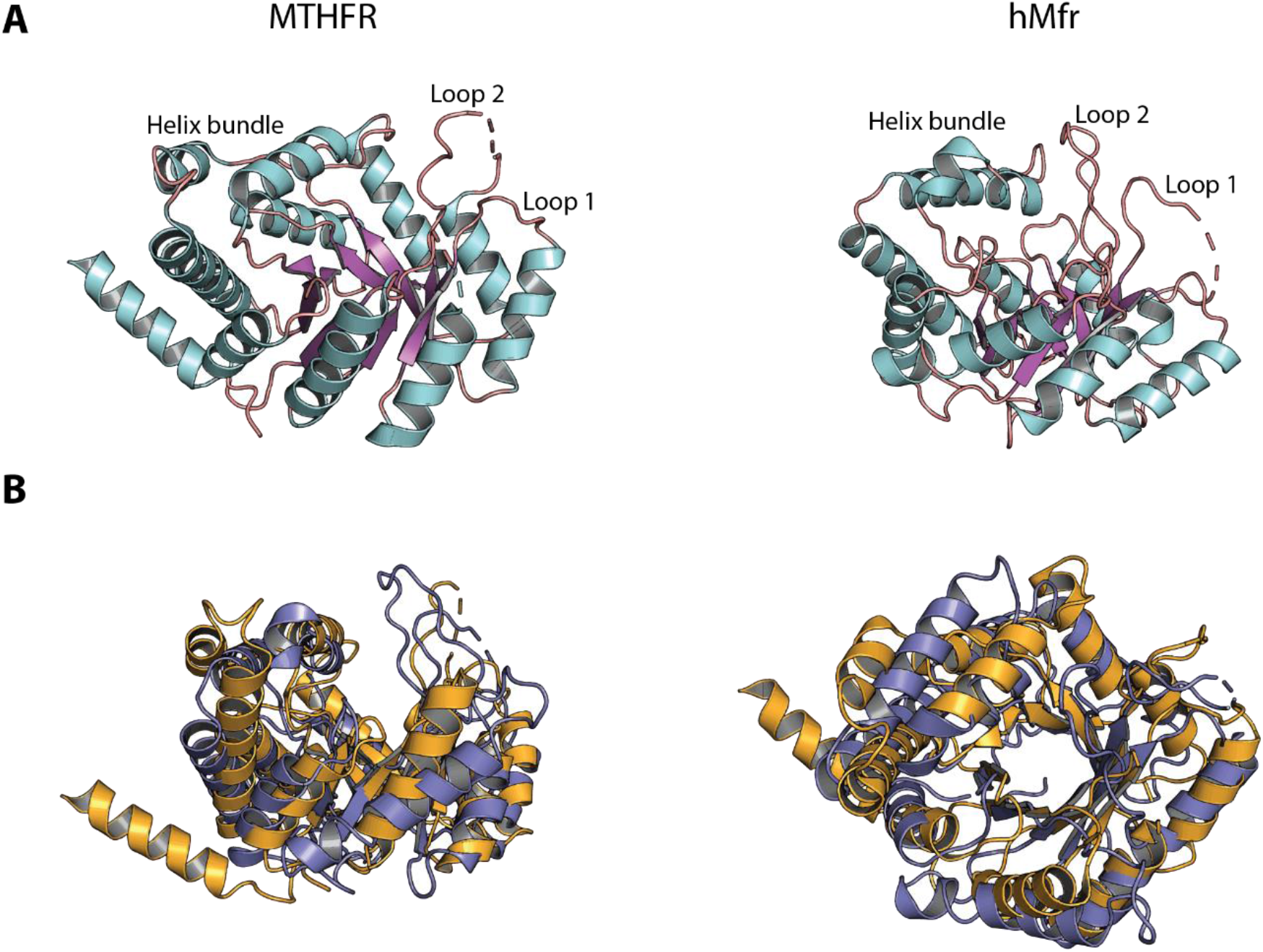
Comparison of the tertiary structures of hMfr (PDB code: 8BGR) and MTHFR from *E. coli.* (PDB code: 1ZP4). (A) Cartoon representation of hMfr and MTHFR with color-coded secondary structure elements. α-Helices are cyan, β-sheets are magenta, and loops are salmon colored. The inserted loops and helix bundle are labeled (see text). (B) The side and top views of the structural alignment of hMfr (blue) and MTHFR from *E. coli* (orange).

**TABLE 3.** X-ray analysis statistics

|  | hMfr |
| --- | --- |
| Data collection |  |
| Wavelength (Å) | 1.0 |
| Space group | *P* 4<sub>3</sub> 2<sub>1</sub> 2 |
| Resolution (Å) | 37.32 - 1.8 (1.9 - 1.8) |
| Cell dimensions |  |
| a, b, c (Å) | 45.79, 45.79, 257.41 |
| $\alpha$, $\beta$, $\gamma$ (°) | 90, 90, 90 |
| R<sub>sym</sub> (%)<sup>a</sup> | 9.3 (111.4) |
| CC<sub>1/2</sub> (%)<sup>a</sup> | 0.996 (0.851) |
| I/σ<sub>I</sub><sup>a</sup> | 9.0 (2.0) |
| Completeness (%)<sup>a</sup> | 94.95 (87.48) |
| Redundancy<sup>a</sup> | 4.3 (3.4) |
| Number of unique reflections | 25392 (2248) |
| Refinement |  |
| Resolution (Å) | 1864-1.8 |
| Number of reflections | 25373 |
| R<sub>work</sub>/R<sub>free</sub><sup>u</sup> (%) | 22.6/24.8 |
| Number of non-hydrogen atoms | 2487 |
| Protein | 2279 |
| Ligands/ions | 0 |
| Solvent | 208 |
| Mean B-value (Å<sup>2</sup>) | 36.38 |
| Molprobity clash score, all atoms | 1.55 |
| Ramachandran plot |  |
| Favoured regions (%) | 99.65 |
| Outlier regions (%) | 0 |
| rmsd<sup>~</sup> bond lengths (Å) | 0.003 |
| rmsd<sup>c</sup> bond angles (°) | 0.59 |
| PDB code | 8BGR |

<sup>a</sup> Values relative to the highest resolution shell are within parentheses. <sup>b</sup> R<sub>free</sub> was calculated as the R<sub>work</sub> for 5% of the reflections that were not included in the refinement. <sup>c</sup> rmsd, root mean square deviation.

### 2.3 Comparison between the structure of hMfr and MTHFR

Both, hMfr and MTHFR are TIM barrel structures (Figure 4B) with a root-mean-square deviation (rmsd) between hMfr and MTHFR from *E. coli* of 4.9 Å in the whole structure region. Unexpectedly, the insertion regions of hMfr and MTHFR, forming the active-site cleft, are architecturally also rather related (see Figure 4B). The binding positions of FAD and methyl-H_4_F were observed in the substrate-containing crystal structure of MTHFR from *E. coli.* The high structural similarity between hMfr and MTHFR from *E. coli* suggests that the equivalent cleft of hMfr could also accommodate the substrates methylene/methyl-H_4_F and NADH/NAD^+^. A multiple sequence alignment of all biochemically characterized Mfrs and MTHFRs with known structure indicates that most of the residues interacting with methyl-H_4_F in MTHFR are conserved in Mfr (Figure 5). While the loop after β2 shields the binding site of both substrates, the prolonged loop segments after β4 and β5 are involved in forming the NAD binding site and those after β7 and β8 are involved in forming the H_4_F binding site. In MTHFR and hMfr, the loop following β4 is partially disordered. A multiple sequence alignment of Mfr family members indicates a high conservation of the insertion segments at the predicted active site (Figure 5).

**Figure 5:**
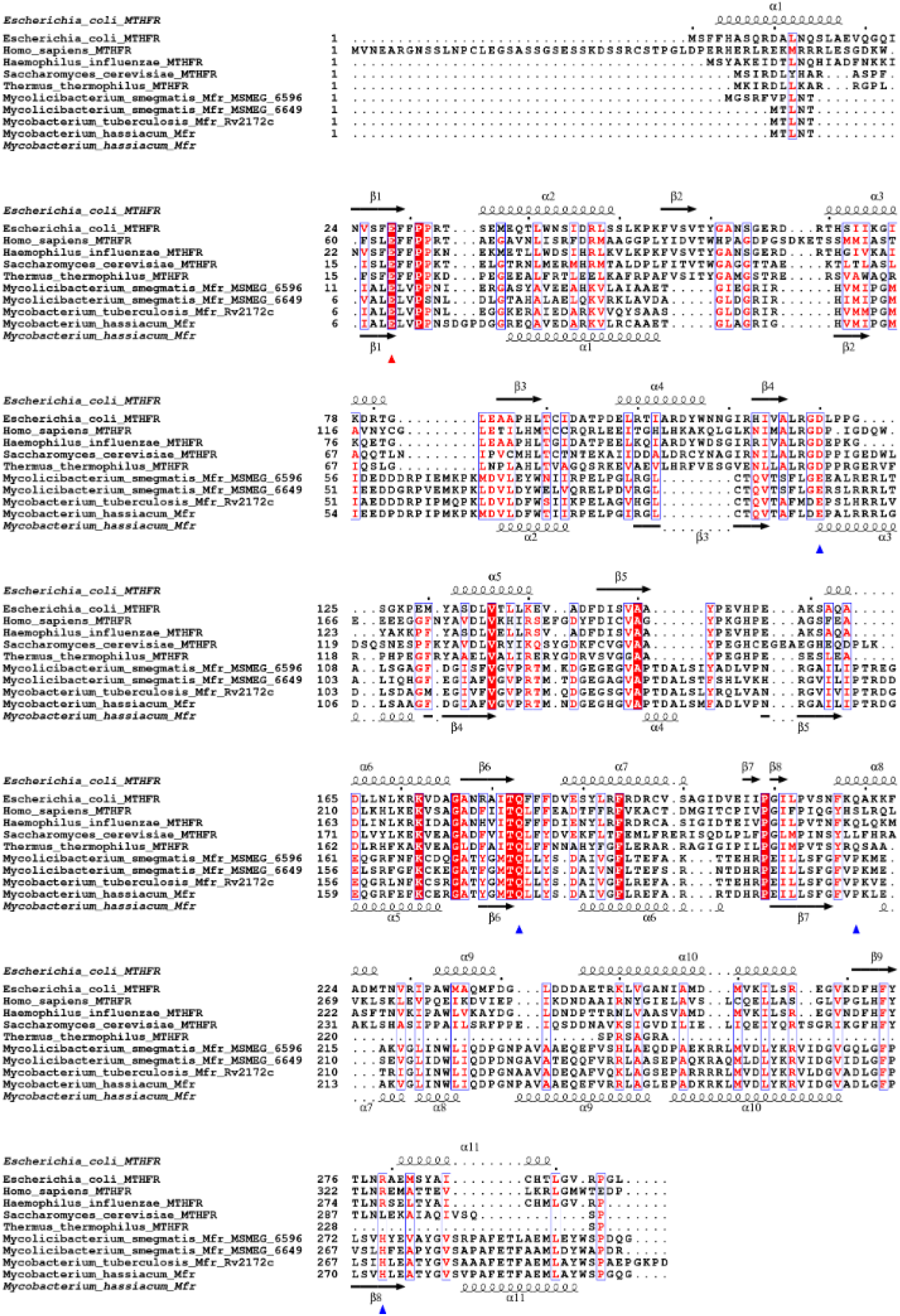
Primary structure comparison of Mfr and MTHFR proteins. The secondary structure elements above the multiple sequence alignment refer to the crystal structure of MTHFR from *E. coli* (PDB: 1ZP4). The secondary structure elements below the alignment refer to the crystal structure of hMfr. Strictly conserved residues are depicted as white letters with red back-ground, residues with similar physicochemical properties are depicted as red letters in blue-lined boxes. The strictly conserved and catalytically essential glutamate residue is marked with a red triangle below the corresponding positon. Residues that directly interact via hydrogen bonds in the ternary complex structure of MTHFR are marked with blue triangles below the corresponding position. For the sake of readability, only the catalytic domains of MTHFR from *Homo sapiens* (residues 1-343) and *Saccharomyces cerevisiae* (residues 1-302) were included. The alignment was generated using MUSCLE,^36^ and the figure was created using ESPript 3.0.^37^

As a consequence, a structural alignment with the crystal structures of hMfr and MTHFR from *E. coli* harboring NADH and methyl-H_4_F supports that the possible binding sites of H_4_F and NAD are located in the aforementioned cleft (Figure 6). The pterin ring of H_4_F confidently accommodates into the predicted pocket of hMfr. The moieties after the phenyl ring clashes with the elongated segment after strand β8, but in the real structure, the flexible tail region may have a different conformation (Figure 6C). The loop 1 (51-69) narrowed the cleft, which is in agreement with an optimized binding of the smaller nicotinamide ring of NADH in comparison with the bulky isoalloxazine of MTHFR (Figure 6C). The adenine dinucleotide moiety of NADH and FAD could be bound to structurally similar sites of the cleft in both enzymes.

**Figure 6:**
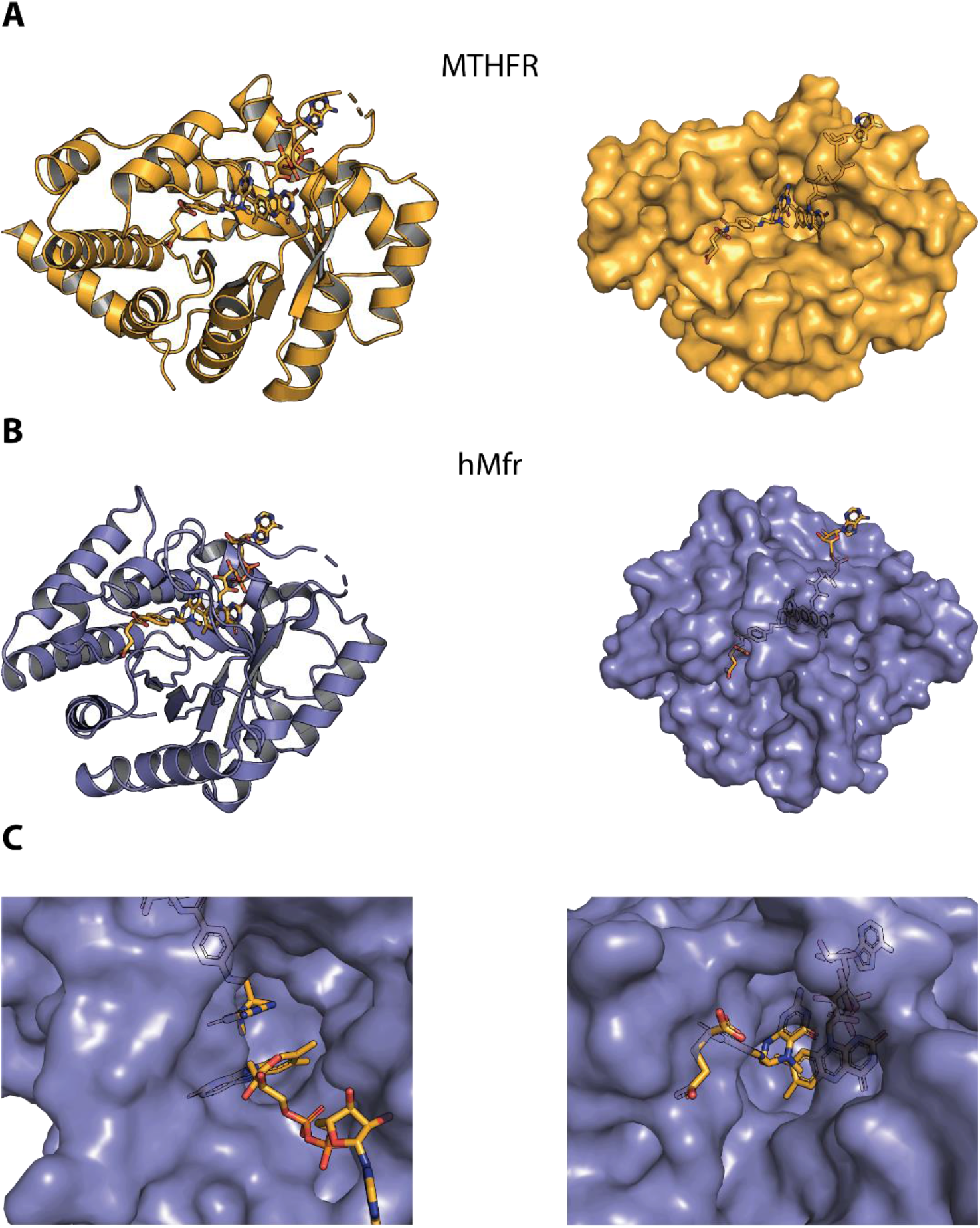
Predicted substrate binding sites of hMfr. (A) Cartoon and surface models of the complex structure of E28Q-MTHFR from *E. coli* in complex with methyl-H_4_F and FAD (PDB code: 1ZP4). The transparency of the surface is 10%, so atoms located behind the image plane can also be shown. (B) Cartoon and surface models of the structure of hMfr. Methyl-H_4_F and FAD from the E28Q-MTHFR complex are superimposed and represented by orange colored carbon atoms. (C) Zoom-up views of the model shown in panel B, showing the protein environment of the pterin ring of methyl-H_4_F and FAD.

### 2.4 Possible active site residues of hMfr

Mutation experiments on MTHFR from *E. coli* revealed several residues important for the catalytic activity.^7–9, 11^ Glu28 is considered as a proton donor for the formation of an iminium cation intermediate, which accepts subsequently the hydride from FADH2. The exchange of Glu28Gln renders the enzyme inactive and led to the only known ternary complex structure of MTHFR. Gln183 is in contact with the pterin ring N1 of methyl-H_4_F via a hydrogen bond. Asp120 interacts with the pterin ring N3 of methyl-H_4_F via a hydrogen bond at a distance of 2.9 Å. Phe223 interacts by π-π interaction with the phenyl ring of methyl-H_4_F in MTHFR.

In the hMfr structure, Glu9 and Gln177 are found in the equivalent positions of Glu28 and Gln183 of MTHFR, respectively (Figure 7). Furthermore, both positions belong to the few amino acids that are conserved in the multiple sequence alignment of Mfrs and MTHFRs (Figure 5). At the position of Asp120 of MTHFR, Glu97 was found in the alignment (Figure 5). However, in the crystal structure of hMfr Glu97 is located at the beginning of α3 and points towards the solvent. Glu55 occupies the position corresponding to Asp120 in the extended loop 1 and probably stabilizes the pterin ring of H_4_F. Leu221 of hMfr is found in the equivalent position of Phe233 of MTHFR, which could stabilize the phenyl ring of H_4_F by hydrophobic interaction (Figure 7).

**Figure 7:**
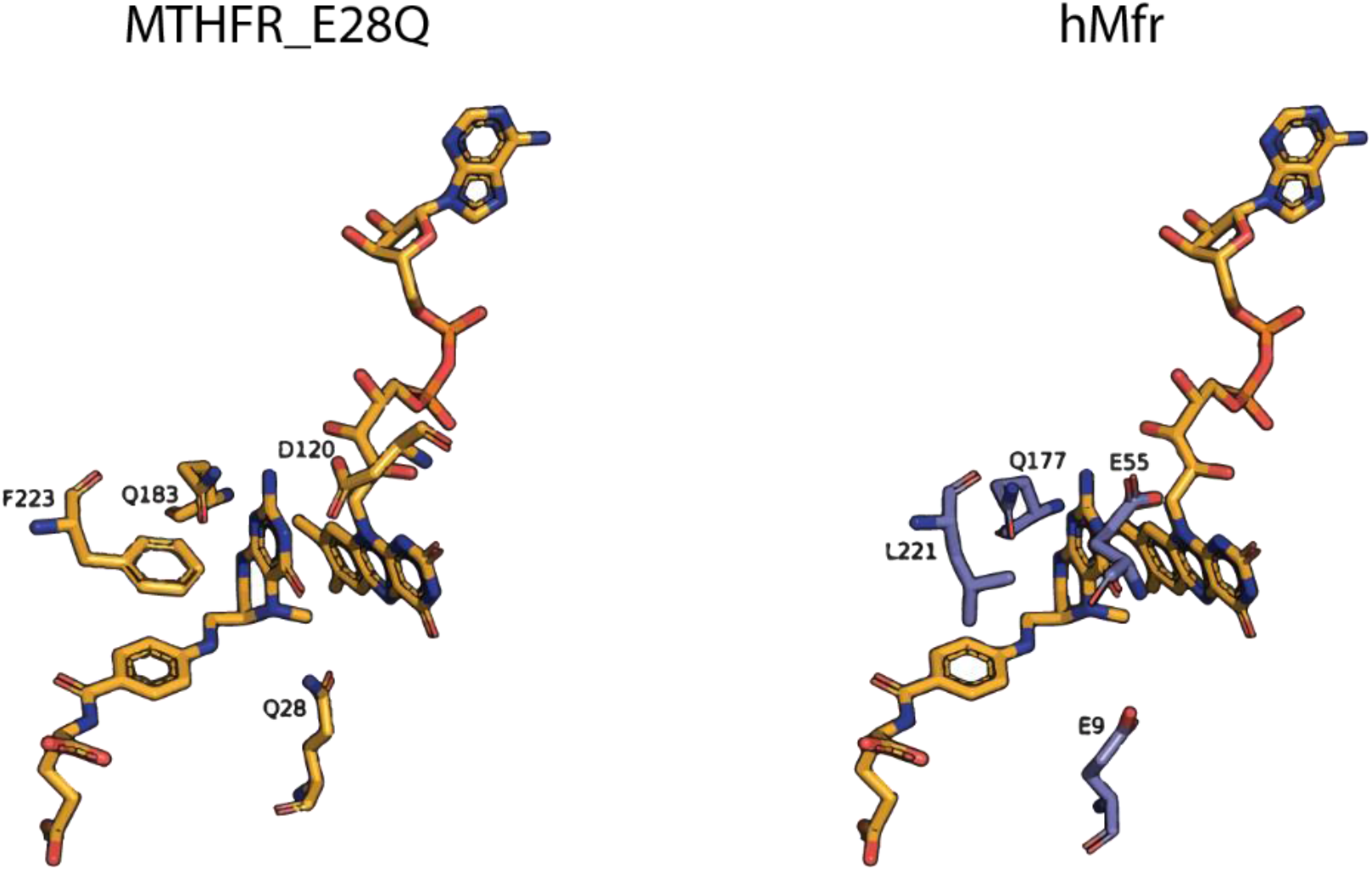
Conservation of the predicted active site residues of hMfr (right) and of the E28Q variant of MTHFR from *E. coli* (left) The residues derived from the crystal structure of the ternary MTHFR complex are shown with orange carbons, while those derived from the structure of the hMfr apoenzyme are shown with light blue carbons.

Based on the structural comparison between MTHFR and hMfr, we hypothesized that methylene/methyl-H_4_F binds to the active site cleft in a highly similar fashion in MTHFR and hMfr as shown in Figure 6 and 7. To substantiate this hypothesis, we constructed an hMfr mutant, in which Glu9 was exchanged to glutamine. The corresponding Glu28Gln mutation of MTHFR from *E. coli* renders the enzyme inactive. As predicted, the catalytic activity of the Glu9Gln variant of hMfr reduced more than 100-fold compared to that of the native enzyme, while the *K_m_* values of the substrates were only slightly changed (Table 2). This finding supported our hypothesis that the geometry of Glu9 relative to methyl-H_4_F is highly similar in hMfr and MTHFR, and that Glu9 protonates the substrate, thereby forming the iminium cation intermediate. This result also implicates that the hydride transfer between C4 of NADH and C14a of methylene-H_4_F occurs at virtually the same site as the hydride transfer from the central pyrazine ring of FAD in MTHFR. The active site geometry for the NADH-to-methylene-H_4_F hydride transfer of hMfr is therefore identical to that of the oxidative half reaction of MTHFR, where a hydride from FADH2 is transferred to methylene-H_4_F.

In the previous studies, Mfr of *M. tuberculosis* was investigated by *in vivo* mutation experiments.^18^ In these experiments, the Arg159Asn and Leu214Ala mutations were introduced into an *M. tuberculosis strain*.^18^ Both strains showed impaired methionine biosynthesis and higher sensitivity to antifolates. Leu214 of Mfr from *M. tuberculosis* is equivalent to Leu221 of hMfr, which is most likely involved in fixing the phenyl ring of H_4_F. Arg159 from *M. tuberculosis* corresponds to Arg148 in hMfr. In the crystal structure, the side chain of Arg148 is far from the possible active site and also does not interact with any of the modeled substrates. However, the side chain is buried in the protein and forms extensive hydrogen bonds to the backbone for stabilization of the structure. Therefore, Arg148 or Arg159 appear to play a structural rather than a catalytic role.

## 3 CONCLUSION

Non-canonical methylene-H_4_F reductase Mfr is a vital enzyme in mycobacteria including the pathogen *M. tuberculosis*. Eukaryotic organisms including humans use instead the different FAD-containing MTHFR for the same reaction. We heterologously overproduced Mfr from the thermophilic *M. hassiacum* and characterized it as a monomeric enzyme that does not contain FAD as prosthetic group. The crystal structure of hMfr revealed a TIM barrel fold, which is also reported for the FAD-binding counterpart MTHFR. Despite the low primary structure similarity and the different enzymatic mechanism, the tertiary structures of hMfr and MTHFR did not only show the same overall fold but also highly related binding sites for methyl-H_4_F. In addition, we mutated Glu9 of hMfr to Gln9, which is equivalent to the Glu28Gln mutant of MTHFR from *E. coli.* This mutation resulted in a strong reduction of the catalytic activity but not in a substantial loss of substrate affinity. NADH appears to occupy the FAD binding site such that the hydride transfer geometry become highly related to that of the oxidative half reaction of MTHFR. The hMfr crystal structure can be applied for designing inhibitors against Mfr, which is an excellent target for antimycobacterial drugs. The unexpected high similarity between hMfr and MTHFRs and conservation of the active site residues in both enzymes suggestes the presence of a common ancestor for both methylene-H_4_F reductases.

## 4 MATERIALS AND METHODS

### 4.1 Construction of the vectors for overproduction of hMfr

The gene sequence of hMfr (Uniprot Accession: K5BDY6) was optimized for the codon usage of *E. coli* and synthesized by GeneScript as described below:

atgaccctgaataccattgcgctggaactggtgccgccgaatagcgatggcccggacggtggtcgtgaacaagcggttgaggatgcgcgtaaggtgctgcgttgcgcggcggagaccggtctggcgggtcgtatcggccacgttatgatcccgggtatgattgaggaagacccggatcgtccgattccgatgaagccgaaaatggacgtgctggatttctggaccatcattcgtccggagctgccgggtatccgtggcctgtgcacccaggttaccgcgtttctggacgaaccggcgctgcgtcgtcgtctgggtgacctgagcgcggcgggttttgatggcattgcgtttgtgggtgttccgcgtaccatgaacgatggtgaaggtcatggtgttgcgccgaccgatgcgctgagcatgttcgcggatctggttccgaaccgtggcgcgatcctgattccgacccgtgacggtgaacagggccgtttcgagtttaagtgcgaacgtggtgcgacctacggcatgacccaactgctgtatagcgacgcgatcgtgggtttcctgcgtgagtttgcgcgtcgtaccgatcaccgtccggaaattctgctgagcttcggttttgtgccgaagctggaagcgaaagttggcctgatcaactggctgattcaagatccgggtaacccggcggtggcggcggagcaagaattcgttcgtcgtctggcgggtctggagccggcggacaagcgtaaactgatggttgatctgtacaaacgtgtgatcgacggtgttgcggatctgggctttccgctgagcgtgcacctggaagcgacctatggtgttagcgtgccggcgtttgaaacctttgcggagatgctggcgtattggagcccgggtcagggttaa

The synthesized gene was cloned into pET-24b(+) between the NdeI and SalI restriction sites. The resulting plasmid pET-24b(+)_hMfr was transformed into chemically competent *E. coli* BL21(DE3) STAR cells (Thermo Fisher Scientific). The recombinant *E. coli* was cultivated at 37 °C overnight in 100 ml of Lysogeny Broth (LB) medium as pre-culture. 20 ml of the pre-culture were used to inoculate two liters of pre-warmed Terrific Broth (TB) medium supplemented with 50 μg/ml kanamycin sulfate in a 2-l flask (Thermo Fisher Scientific). The culture was stirred with 750 rpm and cultivated at 37°C until it reached an OD_600_ of 0.6–0.8. Then the gene expression was induced by the addition of 1 mM (final concentration) of isopropyl-β-D-1-thiogalactopyranoside (IPTG, Sigma-Aldrich). After 3 h of induction at 37 °C, the cells were cooled down in an ice bath and centrifuged for 5 min at 13,000 × g. The cell pellets were stored at −20°C.

### 4.2 Purification of hMfr the *E. coli* cells

All purification steps took place under normal aerobic environment except for the preparation of crystal, the gelfiltration step took place in an anerobic chamber (Koy) filled with a N_2_/H_2_ (95%/5%) atmosphere. Around 3 g of cells were suspended 1:5 in 50 mM Mops/KOH pH 7.0 supplemented with 2 mM dithiothreitol (DTT, Sigma-Aldrich) and disrupted by sonication while in an ice bath using a SONOPULS GM200 (Bandelin) with a KE73 tip with a 50% cycle and 160 W (2 min sonication time with 2 min pauses; in total 5 times). The cell debris were separated by centrifugation at 130,000 × g for 30 min. To separate the thermostable hMfr from the *E. coli* proteins, the cell extract was incubated for 20 min at 50 °C in a water bath and subsequently centrifuged for 20 min at 13,000 × g at 4 °C. Ammonium sulfate (Roth) was added to the supernatant to a final saturation of 40% and the mixture was incubated under constant stirring in an ice bath for 20 min and then centrifuged for 20 min at 13,000 × g. The resulting supernatant was applied to two tandemly connected 5 ml-HiTrap Phenyl Sepharose HP columns (Cytiva) equilibrated with 50 mM Mops/KOH pH 7.0 containing 2 mM DTT and 1.6 M ammonium sulfate. The column was washed with 1.1 M ammonium sulfate in the equilibration buffer. Elution proceeded in a single step with 0.7 M ammonium sulfate in 50 mM Mops/KOH pH 7.0 containing 2 mM DTT. The eluted protein fraction was pooled and desalted with 50 mM Mops/KOH pH 7.0 containing 2 mM DTT using a HiPrep 26/10 desalting column containing Sephadex G-25 resin. The desalted sample was applied to a 6 ml Resource Q column equilibrated with the same buffer. Elution took place with a linear gradient from 0–250 mM NaCl over 15 column volumes. hMfr eluted at a conductivity of around 12 mS/cm. The corresponding fractions were pooled, concentrated with a Amicon filters (10 kDa cutoff, Merck) and applied to a HiPrep 26/60 Sephacryl S-100 HR column equilibrated with 20 mM MOPS/KOH pH 7.0 containing 2 mM DTT, 150 mM NaCl and 5% (v/v) glycerol. hMfr eluted around 60 ml retention volume. The corresponding fractions were collected and used for the preparation of anaerobic crystal plates and the measurement of the UV/vis spectrum of hMfr.

For the preparation of hMfr for enzyme assays, a shorter method was applied. After the heat treatment, the supernatant was diluted 1:1 with 50 mM Mops/KOH pH 7.0 containing 2 mM DTT and the diluted protein solution was directly applied to the Resource Q column and eluted as described above. The corresponding fractions were collected and used for the determination of the optimum temperature and the kinetic constants.

### 4.3 Enzymological assay

The activity assay of hMfr was carried out in 1 ml quartz cuvettes filled with 0.7 ml of 100 mM Mops/NaOH pH 7.0 under an N_2_ atmosphere and incubated at 40 °C. In the standard assay, NADH (Roth) and methylene-H_4_F (Schircks Laboratories) were added to final concentrations of 100 μM and 300 μM, respectively. For the kinetic analysis, the concentration of NADH and methylene-H_4_F was set as the standard condition and the concentration of the counter substrate was varied, as shown in Figure S1. The assay was started by the addition of hMfr. The decrease of NADH absorbance at 340 nm was measured (ε_340_ = 6.22 mM^−1^ cm^−1^). For the determination of the temperature dependency, the standard conditions were used and the cuvettes were incubated at different temperatures as indicated in Figure 3.

### 4.4 Crystallization

Crystallization was performed in an anaerobic chamber filled with a N_2_/H_2_ (95%/5%) gas mixture at 8 °C. For sitting drop vapor diffusion method, 96-well screening plates were filled with 90 μl of reservoir solution and the 48-well optimization plates with 200 μl of reservoir solution. The crystallization plates filled with the reservoir solutions were set up in the chamber one week prior of crystallization to make the solution anaerobic. The first crystals were found in several solutions of JB Screen Classic 1 (Jena Bioscience); for example, 25% polyethylene glycol (PEG) monomethyl ether 2000. During the optimization small rod-shaped crystals and thin plates emerged in several PEG solutions (PEG1000–PEG 8000) in a concentration range between 10–30% (v/v) with no additives and no buffers. The best diffracting crystal was a thin plate, which grew in 15% (v/v) PEG 6000 with a final concentration of 30 mg/ml hMfr in the drop over the course of 4 weeks. The best crystal was transferred into the reservoir solution supplemented with 30% (v/v) glycerol and frozen.

### 4.5 X-ray crystallographic analysis

The diffraction experiments were performed at 100 K on the beamline SLS Beamline X10SA (Villigen, Switzerland) equipped with a Dectris Eiger2 16 M detector.

The dataset was processed with XDS and scaled with XSCALE ^21^.The phase problem was solved by the molecular replacement method using PHASER ^22^ with an Alphafold2 ^23^ model of hMfr. The Alphafold2 model was generated by the Colab Notebook ^24^ integrated into the CCP4 suite ^25^The model was built and improved in in COOT ^26^ and refinement was done using Phenix.refine ^27^ and Refmac ^28^. The final model was validated using the MolProbity ^29^ implement of Phenix. Data collection, refinement statistics and PDB code for the deposited structure are listed in Table 3. The figures of the structures and the ternary complex model were generated using PyMOL (version 2.4.1, Schrödinger).

### 4.6 Kinetic data processing and figure generation

The graphs and the kinetic constants analyses were performed with Python 3.7 using Jupyter Notebook (version 6.1.4) ^30^ as development environment and the following packages: os, pandas ^31^, seaborn ^32^, matplotlib ^33^, numpy ^34^, scipy ^35^.The determination of the kinetic constants is based on the experimentally determined average values of the specific activity and the corresponding standard deviation from three replicates per measurement point. The Michaelis-Menten equation (vi = vmax * substrate/(km + substrate)) was used as the corresponding function for scipy.curve_fit. The resulting standard deviations for the curve fitting process were calculated from the output of the pcov parameter from scipy.cuve_fit by calculating the diagonal of the resulting array and subsequently taking the square root of the diagonal.

## AUTHOR CONTRIBUTIONS

**Manuel Gehl**: Conceptualization (equal); data curation (equal); formal analysis (equal); investigation (equal); methodology (equal); software (equal); validation (equal); writing – original draft (supporting); writing – review and editing (equal). **Ulrike Demmer:** data curation (supporting); investigation (supporting). **Ulrich Ermler**: Conceptualization (supporting); funding acquisition (equal); project administration (equal); resources (equal); data curation (equal); formal analysis (equal); investigation (supporting); methodology (equal); software (equal); validation (equal); writing – review and editing (equal). **Seigo Shima**: Conceptualization (equal); funding acquisition (equal); project administration (equal); resources (equal); supervision (equal); writing – original draft (equal); writing – review and editing (equal).

## ACKNOWLEDGEMENT

M.G. thanks the International Max Plank Research School thesis-committee members: Tobias Erb and Rolf Thauer.

## DATA AVAILABILITY STATEMENT

Data available in article supplementary material.

## SUPPORTING INFORMATION

Additional supporting information can be found online in the Supporting Information section at the end of this article.

## SUPPORTING INFORMATION

**Table S1:** Raw data measured for the kinetic evaluation of Mfr and Mfr_E9Q.

| Mfr_Wild type |  |  |  |
| --- | --- | --- | --- |
| Concentration NADH [μM] | Measurement 1<br>[U/mg] | Measurement 2<br>[U/mg] | Measurement 3<br>[U/mg] |
| 0 | 0.00 | 0.00 | 0.00 |
| 20 | 10.07 | 7.40 | 9.77 |
| 40 | 12.14 | 10.66 | 11.85 |
| 60 | 13.50 | 14.63 | 14.63 |
| 80 | 12.13 | 14.52 | 13.97 |
| 100 | 13.24 | 14.08 | 14.55 |
| 130 | 14.63 | 14.41 | 15.31 |
| Concentration CH2=H4F [μM] | Measurement 1<br>[U/mg] | Measurement 2<br>[U/mg] | Measurement 3<br>[U/mg] |
| 0 | 0.00 | 0.00 | 0.00 |
| 50 | 5.03 | 3.26 | 3.55 |
| 100 | 5.63 | 7.11 | 6.52 |
| 200 | 8.88 | 9.77 | 9.77 |
| 300 | 13.24 | 14.08 | 14.55 |
| 400 | 12.14 | 12.14 | 14.51 |
| 500 | 13.33 | 11.55 | 12.14 |
| Mfr_E9Q |  |  |  |
| Concentration NADH [μM] | Measurement 1<br>[U/mg] | Measurement 2<br>[U/mg] | Measurement 3<br>[U/mg] |
| 0 | 0.000 | 0.000 | 0.000 |
| 25 | 0.010 | 0.008 | 0.010 |
| 30 | 0.016 | 0.012 | 0.017 |
| 40 | 0.019 | 0.021 | 0.017 |
| 75 | 0.026 | 0.023 | 0.021 |
| 95 | 0.022 | 0.022 | 0.031 |
| 150 | 0.025 | 0.026 | 0.027 |
| Concentration CH2=H4F [μM] | Measurement 1<br>[U/mg] | Measurement 2<br>[U/mg] | Measurement 3<br>[U/mg] |
| 0 | 0.000 | 0.000 | 0.000 |
| 20 | 0.003 | 0.002 | 0.005 |
| 50 | 0.005 | 0.007 | 0.007 |
| 100 | 0.010 | 0.009 | 0.010 |
| 200 | 0.019 | 0.018 | 0.019 |
| 300 | 0.023 | 0.028 | 0.020 |
| 500 | 0.025 | 0.029 | 0.023 |

**Figure S1:**
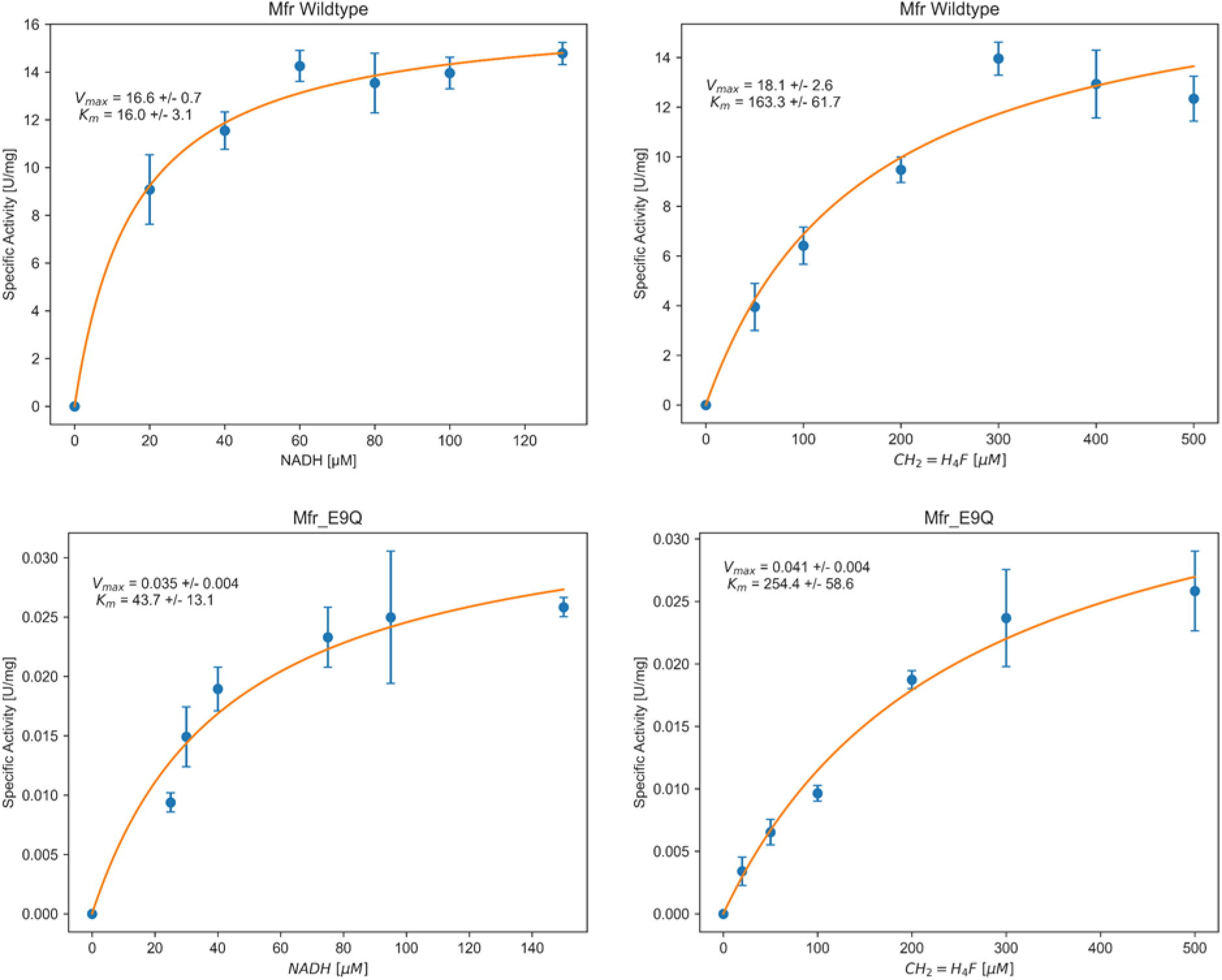
Kinetic fitting for the measurements of hMfr wild type and hMfr_E9Q.

**Figure S2:**
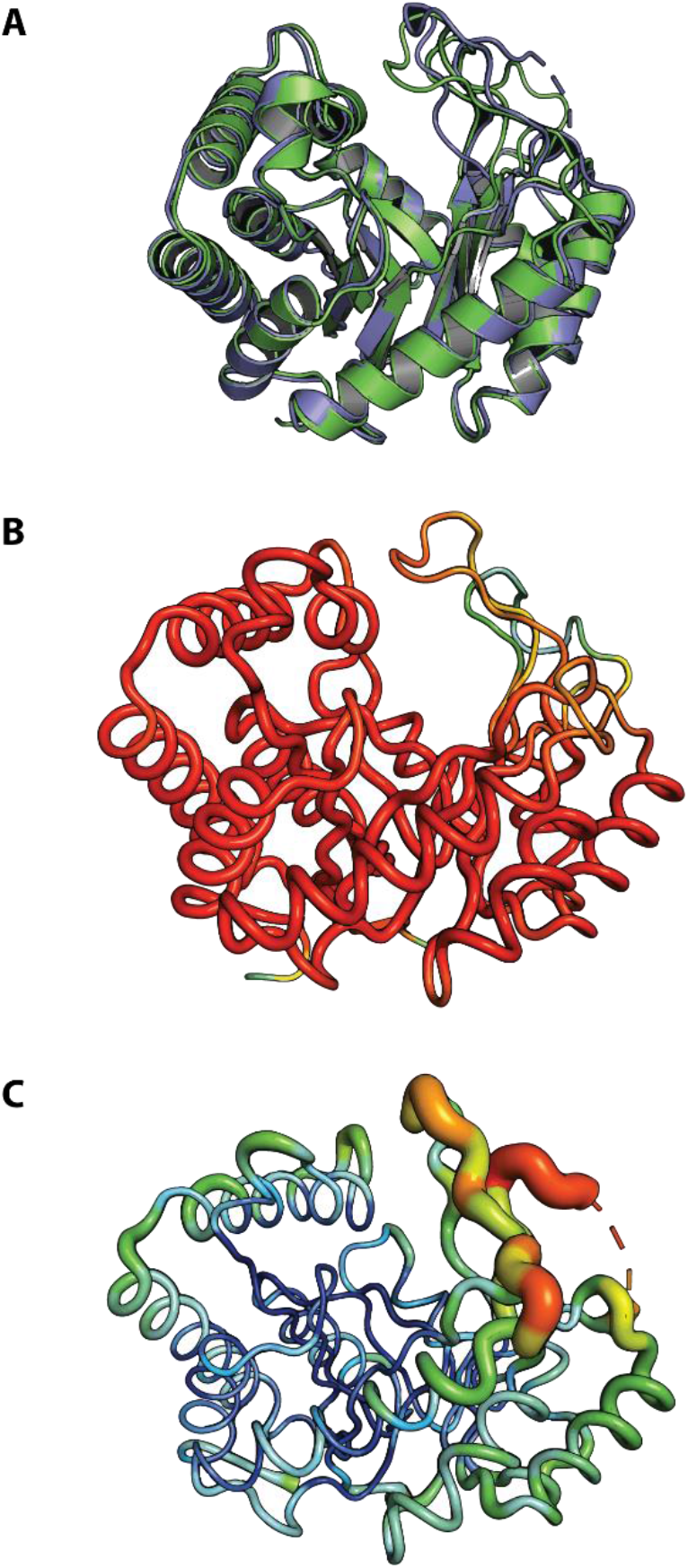
(A) Crystal structure of hMfr (light blue) and the Alphafold2 model (green), which is used as the template for molecular replacement. The overall RMSD between all amino acids is 0.51. (B): Alphafold2 model of hMfr depicted according to the parameter for local distance difference test (pLDDT) values. Red colors indicate highest value of pLDDT with high prediction confidence, while blue colors indicate lowest values of pLDDT with low prediction confidence of the model (C) Atomic displacement parameter (ADP) value of the crystal structure of hMfr. Blue colors indicate a low ADP value, while red colors indicate a high ADP value.

